# GAL-101 prevents amyloid beta-induced membrane depolarization in two different types of retinal cells

**DOI:** 10.1101/2025.01.21.634046

**Authors:** Erika Pizzi, Simona Gornati, Stefano Stabilini, Dario Brambilla, Chiara A. Mercurio, Hermann Russ, Christopher G. Parsons, Michele Mazzanti

**Affiliations:** Dept. Bioscienze, University of Milan, Via Celoria 26, 20133 Milano, Italy; Dept. Fisiopatologia e dei Trapianti, University of Milan, Via Mangiagalli 32, 20133 Milano, Italy; Galimedix Therapeutics Inc. Kensington, MD 20895, USA

## Abstract

Glaucoma and age-related macular degeneration (AMD) are two of the major causes of progressive vision loss and ultimately blindness worldwide. Both retinopathies share several pathological features with Alzheimer’s disease (AD) such as: impairment of neuronal function, astrocytosis, and activation of immune-competent microglia and Müller cells. It also has been shown that these conditions are characterized by the presence of an elevated concentration of amyloid beta (Aβ). Under pathological conditions, Aβ_1-42_ tends to aggregate, forming toxic soluble oligomers, considered to be the most harmful amyloid species. One strategy adopted to prevent cell damage caused by these oligomers is to impair their aggregation. Here we studied GAL-101, a small molecule designed to modify the aggregation of Aβ_1-42._ To assess the role of GAL-101 in the aggregation of Aβ_1-42_, *in vitro* electrophysiological measurements on retinal ganglion cells (RGCs) and retinal pigment epithelial (RPE) cells were performed to determine the polarization of the resting membrane potential. Cells treated only with Aβ_1-42_ oligomers showed a strong depolarization of the resting membrane potential, which is believed to be the main reason for retinal cells malfunctioning in neurodegenerative diseases of the eye. Pre-incubation with GAL-101 stabilized the cell resting potential to around -50mV during exposure to Aβ_1-42_, in both RGCs and RPE cells. GAL-101 was able to prevent changes in resting membrane potential and thus would be expected to prevent impairment of retinal cell function. These results are supportive of evaluating GAL-101 as a potential treatment of Aβ-associated retinopathies like glaucoma and dry AMD.

## 1. Introduction

Glaucoma and age-related macular degeneration (AMD) are leading causes of progressive vision loss and blindness worldwide, and their incidence and prevalence increase with age (1–3). While recognized as distinct pathologies, similarities exist between these two retinal diseases, as well as with Alzheimer’s disease (AD). AD is a neurodegenerative disease characterized histopathologically by the presence of extracellular deposits of misfolded amyloid-beta (Aβ) in the form of senile plaques and the intracellular accumulation of hyperphosphorylated tau in the form of neurofibrillary tangles (4). The incidence of all three conditions increases with age and the chronic neurodegenerative changes seen in the eyes of glaucoma and AMD patients are similar to changes characteristic of the brains of AD patients (1–3,5–7).

The association between glaucoma and AD has emerged from studies showing that AD is associated with glaucomatous changes, such as RGC loss, optic neuropathy and impaired visual function (8–13). Aβ has been reported to be implicated in the development of RGC apoptosis in glaucoma, with evidence of caspase-3-mediated abnormal APP processing (14), increased expression of Aβ_1-42_ in RGCs and optic nerves (14–19) and decreased vitreous Aβ levels (consistent with retinal Aβ deposits) in patients with glaucoma (20).

AMD is characterized by drusen, extracellular waste deposited between the basal surface of the retinal pigment epithelium (RPE) and Bruch’s membrane (21). Drusen are a clear hallmark of early, or “dry” AMD (22), and a significant risk factor for progression to “wet” AMD involving neovascularization (23). Drusen include deposits of Aβ, remnants of RPE cells (24), and a variety of immune system-related molecules, including immunoglobulins and complement system components (25). The deposition of Aβ, in particular Aβ oligomers, in drusen has been shown to increase with age (26,27). Small soluble oligomeric forms of Aβ are able to induce significant RGC apoptosis *in vivo* and *in vitro* (16,28,29). It has been shown that targeting Aβ_1-42_ formation and aggregation reduces glaucomatous RGC apoptosis *in vivo* and therefore raises the possibility of using antiaggregant therapy to provide neuroprotection against glaucoma progression (16). GAL-101 (formerly MRZ-99030, EG30) is a small D-amino acid dipeptide that modulates Aβ_1-42_ aggregation by triggering a non-amyloidogenic aggregation pathway and thereby reduces the amount of toxic soluble oligomeric Aβ_1-42_ species (30–32).

According to recent publications, Aβ is able to form ionic pathways of different conductance depending on the size of the oligomeric species (33–36). This might be related to the ability of Aβ to infiltrate biological membranes, as has been described in detail, into an artificial lipid bilayer (37,38).

In the present study, we monitored the resting membrane potential (RMP) of RGC and RPE cells before and after acute application of 50nM Aβ_1−42_. The gating of unspecific ionic pathways mediated by Aβ_1−42_ caused a net inward current, thus inducing RMP depolarization. Preincubation of Aβ_1-42_ with GAL-101 was able to reduce or even prevent RMP depolarization.

## 2. Materials and Methods 2.1.Cell culture

RGCs were isolated from P5-P6 mouse retinas (neonatal mice C57BL/6N, strain code 027, Charles River) using the standardized procedure in the Miltenyi Biotec kit. By the 4th day *in vitro*, three types of cells were distinguishable: RGCs, amacrine cells, and astrocytes. RGCs were recognized by the presence of one long protrusion representing the axon. RGCs were maintained for 7 days in Neurobasal medium supplemented with Sato 1X, B27 2X, T3 1X, Sodium Pyruvate 1 mM, Glutamine 1 mM, Insulin 1X, CNTF 1X, BDNF 1X and Forskolin 1X at 37°C and 5% CO2 before recording.

RPE cells were obtained from the American Tissue Collection (ARPE-19 - ATCC^®^ CRL-2302TM) and maintained in 1:1 DMEM/F12 with 10% FBS at 37°C and 5% CO_2_.

### 2.2. GAL-101 molecule

GAL-101 (see Figure 1) is designed to prevent the formation of all forms of toxic Aβ oligomers by binding with high affinity to the misfolded Aβ monomers before they can form toxic soluble oligomers. These oligomers then rapidly form amorphous, non-beta-sheet “clusters,” which are innocuous. Interestingly, once GAL-101 concentration reaches effective levels, it “triggers” formation of the “clusters”, which have shown their ability to collect additional misfolded Aβ monomers, even in the absence of additional GAL-101 molecules, through a self-propagating mechanism. This novel “trigger effect,” protected by Galimedix’s patent portfolio, results in a sustained effect. The effect lasts far longer than the time a single administration of the drug remains at therapeutic levels in the retina, potentially allowing for a convenient interval application regimen for patients. Thus, GAL-101 drops may potentially provide sustained prevention of formation of toxic Aβ oligomers in the retina, leading to a reduction in complement response and its consequent damage. Thus GAL-101 could contribute to slowing or stopping progression, and possibly restoring neural function depressed by the chronic toxic attack.

**Figure 1:**
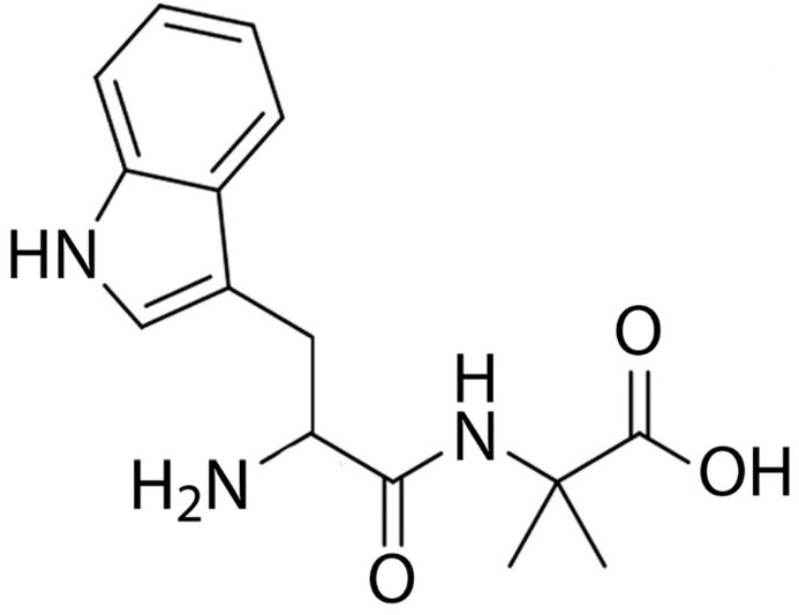
Chemical structure of GAL-101

### 2.3. Electrophysiological recordings

#### 2.3.1. Resting membrane potential measurements

Membrane potential recordings were conducted on single cells using the perforated whole-cell configuration in current-clamp mode. Data were collected using an Axopatch 200B amplifier (Molecular Device, CA, US) and experimental traces were digitized at 5 kHz and filtered at 1 kHz with a Digidata 1322 acquisition system. Clampex-8 was used as the acquisition software. Patch pipettes (GB150F-8P with filament, Science Products) were pulled from hard borosilicate glass on a Brown-Flaming P-87 puller (Sutter Instruments, Novato, CA, US) and fire-polished to a final electrical resistance of 4-5 MΩ. Shortly before starting the recordings, the culture medium was replaced with the external recording solution. The perforated whole-cell configuration was achieved by using the antibiotic Gramicidin (Sigma Aldrich) diluted in the internal solution at a final concentration of 5 μg/ml and 10 μg/ml for RGCs and RPEs, respectively. Electrical access to the cell was thereby achieved after about 5-10 minutes.

The solutions used were (in mM):

RGCs –solution ext.: 140 NaCl, 2.5 KCl, 1.8 CaCl_2_, 0.5 MgCl_2_, 10 Glucose, 10 HEPES, pH 7.4.

RGCs - solution int.: 140 KCl, 10 NaCl, 2 MgCl_2_, 0.1 CaCl_2_, 10 Glucose, 10 HEPES, pH 6.9.

RPE cells – external solution: 132 NaCl, 2 CaCl_2_, 1 MgCl_2_, 10 Glucose, 10 HEPES, pH 7.4.

RPE cells - internal solution: 132 KCl, 2 CaCl_2_, 1 MgCl_2_, 10 Glucose, 10 HEPES, pH 7.0.

#### 2.3.2. HEK293 whole-cell voltage-clamp recordings

Patch-clamp electrophysiology was performed in whole-cell configuration in HEK293 cells. The intracellular solution was (in mM): 20 KCl, 120 K-aspartate, 10 HEPES, and 1 MgCl_2_. The potassium channel blocker tetraethyl-ammonium (TEA 40 mM; Sigma, St Louis, MO, USA) was used to minimize the contribution of these ions. To obtain voltage-current relationships (i/V), the cell voltage was held at -50 mV and the current was measured at the end of 800 ms voltage steps from -70 to + 50 mV. A subtraction method using 50 µM indanyloxyacetic acid 94 (IAA94, Sigma, St Louis, MO, USA) a Cl-channel inhibitor, was used to isolate the inhibitor-sensitive current. The Axopatch 200 B amplifier and the pClamp-9 acquisition software and Clampfit 9 (both from Molecular Device, Novato, CA) were used to record and analyze whole-cell traces. Current recordings were digitized at 5 kHz and filtered at 1000 Hz. Patch electrodes (GB150F-8P with filament, Science Products) were pulled from hard borosilicate glass on a Brown-Flaming P-87 puller (Sutter Instruments, Novato, USA) and fire-polished to a tip diameter of 1-1.5 µm and electrical resistance of 5-7 MΩ.

#### 2.3.3. Tip-dip lipid bilayer current recording

Single-channel recordings from lipid bilayers were obtained using the tip-dip method. In brief, patch clamp pipettes (Garner Glass 7052) were made using a P97 Sutter Instruments puller (Novato, CA) and fire-polished to a tip diameter of 1–1.5 μm and 5–7 MΩ resistance. The same solution was used both in the bath and in the pipette (140 mM KCl, 10 mM Hepes, pH 6). Purified recombinant CLIC1 protein (2 μg/ml) was added to the pipette solution. As soon as the pipette tip reached the bath solution, a phospholipid monolayer (phosphatidylcholine, Avanti Polar Lipids, Inc., Birmingham, AL) was spread on the surface. The electrode was repeatedly passed through the surface of the solution until the pipette resistance rose above 5 GΩ. An Axopatch 1D amplifier and pClamp 7 (both from Axon Instruments, Novato, CA) were used to record and analyze single-channel currents. Current recordings were digitized at 5 kHz and filtered at 800 Hz.

### 2.4. Amyloid beta preparation

Aβ was prepared according to the protocol originally developed by the group of Klein (32,39). In brief: lyophilized Aβ_1-42_ from Bachem (Code. H-1368 lot number 1030255) was dissolved in HFIP (Sigma Aldrich) at a concentration of 1 mg/ml under 90 min of continuous shaking. 50 ml aliquots were then frozen at –80 °C for 45 minutes. After overnight lyophilization (−20 °C) the aliquots were kept stored at –20 °C until use.

Right before use, Aβ_1-42_ was dissolved in DMSO at a final concentration of 100 μM and sonicated in an ultrasonic water bath for 1 hour. Aβ_1-42_ was quantified by using a BCA protein assay kit (Pierce). DMSO concentration never exceeded 1% (ranging from 0.1% to 0.5%).

No previous data have been reported in the literature on the use of Aβ_1-42_ on single isolated RGCs or RPE cells. We therefore designed our experiment based on our own preliminary experiments. The final Aβ_1-42_ concentration was different depending on the cell type. The concentration-response analysis revealed the optimal concentration for RGCs to be 50 nM. In contrast, the equivalent concentration for RPE cells was 1 μM as RPE cells were considerably less sensitive.

In all experiments, Aβ_1-42_ was used after a 90-minute incubation period at 36 °C and within 2 hours from its dilution in the external solution. Aβ_1-42_ was applied using a gravity-driven perfusion system (RSC-200, BioLogic) through a micropipette positioned close to the recorded cell.

GAL-101 was used at the final concentration of 0.5 and 1 μM for RGCs and 10 and 20 μM for RPE cells, corresponding to ten or twenty times of the Aβ_1-42_ concentration. GAL-101 was added to Aβ_1-42_ before incubation.

### 2.5. Voltage-clamp, whole-cell configuration

Aβ_1-42_ was incubated at 37 °C at different times at the concentration of 1 µM (see methods). HEK cells were exposed to different preparations of Aβ_1-42_ for 15 minutes prior to the measurement of membrane conductance. The final Aβ_1-42_ concentration in contact with cells was 50 nM.

### 2.6. Statistical analysis

The statistical significance of the data was assessed using a two-tailed paired Student’s t test. All data are shown as mean ± SEM. Origin 9 was used for plotting data and for statistical analysis.

## 3. Results

### 3.1. Amyloid beta impact on resting membrane potential of retinal ganglion cells

To evaluate the impact of Aβ_1-42_ on the RMP of RGCs, we first measured the resting potential of the cells after performing whole-cell patch-clamp recordings in current-clamp mode. Under controlled conditions, RGCs had an average RMP of -68.3 ± 1.9 mV (n=26; Figure 2A). The RMP of RGCs also was measured under control conditions up to 1 hour (Figure 2B) to be sure that our experimental recording conditions were not affecting cell functions. In some cells, it was not unusual to observe spontaneous firing activity (Figure 2C).

**Figure 2:**
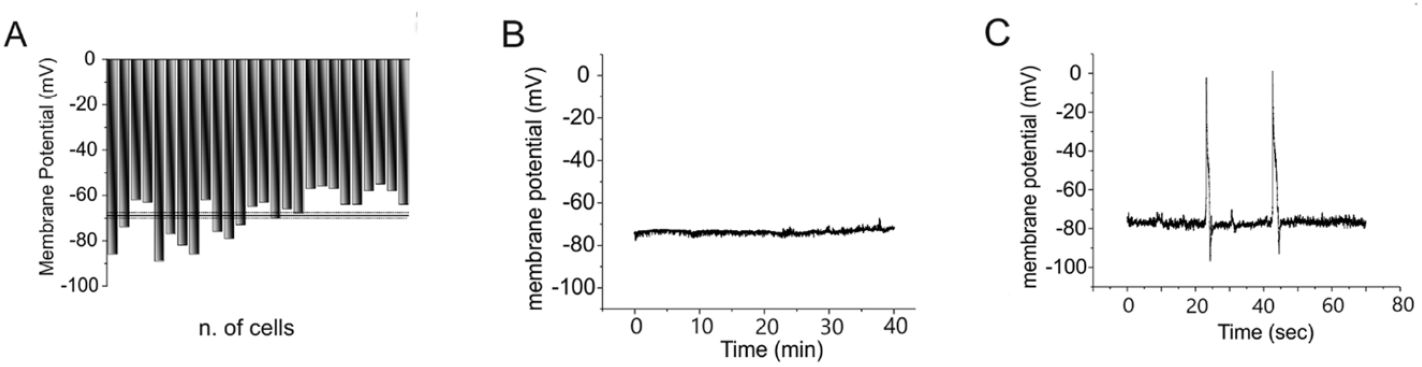
Measuring resting potential in RGC-isolated cell. A: Distribution of cell resting potential (average -68.3 ±1.9mV;n = 26). B: Stability of membrane potential over 40 minutes of continuous recording (n = 12). C: Spontaneous action potential in RGC cell during resting membrane potential monitoring.

The Aβ_1-42_ concentration was able to induce a relevant membrane voltage depolarization without being toxic for the cell. This was estimated by performing a dose/response curve to calculate the EC50 (half maximal effective concentration) of Aβ_1-42_. To determine this, we used three different procedures; Figure 3 shows two of the three. The first method (Figure 3A) consisted in measuring the RMP in time course experiments under controlled conditions for several minutes. Aβ_1-42_ was applied to the cells from 1 nM up to 10 nM Aβ_1-42_, with 1 nM increments every 4 minutes, and from 50 to 100 nM. Using the second method (Figure 3B), the Aβ_1-42_ effect on RMP was measured using a peptide concentration from 10 nM up to 100 nM delivered every 4 minutes. Average RMP values at each concentration (n = 3) obtained by these two experimental procedures were used to generate the dose-response curve shown in Figure 3C. The plot depicts combined data from the two protocols previously described (squares) as well as from a third method in which Aβ_1-42_ was applied on a current-clamped RGC as a single concentration (n = 3; circles). From the curve fitting, we obtained an EC50 of 44.06 ± 6.3 nM (membrane potential depolarization -35.6 ± 1.8 mV) and a maximal effect at 100 nM. According to this result, all the experiments on RGCs with GAL-101 were performed using 50 nM Aβ_1-42_, an effective concentration to induce a membrane potential depolarization without causing short term toxicity.

**Figure 3:**
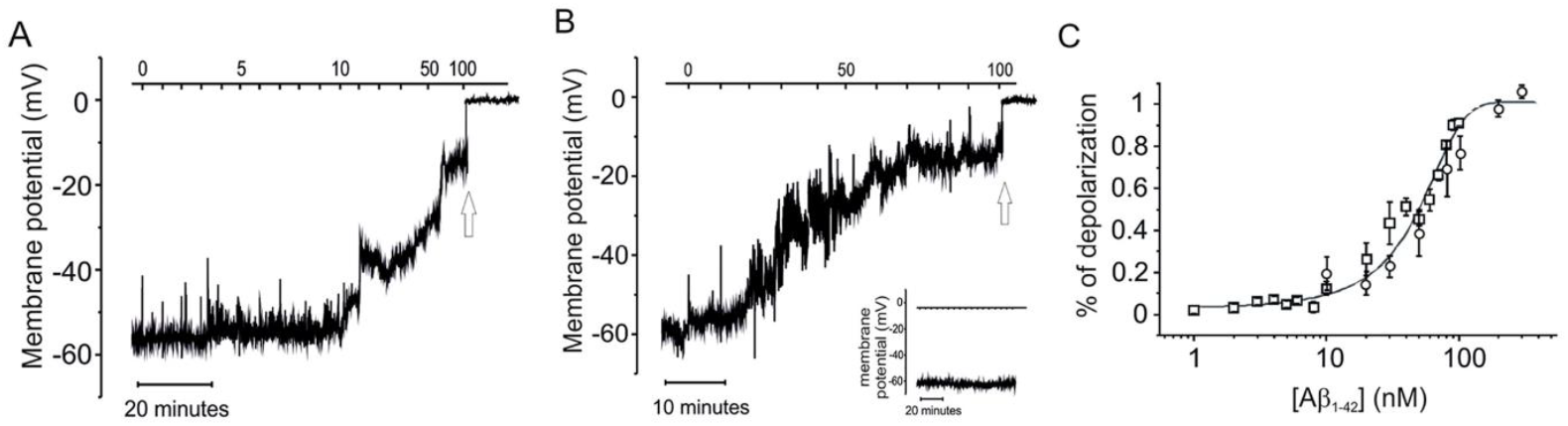
Time course of whole-cell current-clamp experiments. A and B: Depolarization ofRGC cells after addition of Aβ_1-42_ at increasing concentrations (A:n = 3 B:n = 3). The insert in B shows control measurement of a stable membrane potential over 80 minutes. C: Dose/response curve of depolarization rate, recorded in a dynamic condition (circles, n = 3) and from steady-state experiments (squares, n = 3). Calculated EC50 is 44.1 ±6.3 nM at a membrane voltage of-35.6 ± 1.8 m V.

### 3.2. GAL-101 prevents beta amyloid aggregation and reduces membrane depolarization in RGCs

We have previously demonstrated that Aβ_1-42_ aggregation into oligomers increases the permeability to cations in biological membranes (35). Since GAL-101 is a modulator of Aβ_1-42_ oligomerization, we assessed whether it is able to prevent the depolarization of the RMP induced by oligomeric Aβ_1-42_ in RGCs. To this end, the RMP of isolated RGCs was monitored under whole-cell, current-clamp conditions and perfused with a solution containing 50 nM Aβ_1-42_ preincubated with different concentrations of GAL-101. As previously shown, Aβ_1-42_ is already effective in causing a membrane potential depolarization at 20 nM. As shown in Figure 4, GAL-101 inhibited Aβ_1-42_ -induced depolarization in a concentration-dependent manner. Indeed, the GAL-101 effect was statistically significant at a 20-fold (1 µM) stoichiometric increase over Aβ_1-42_ compared to 0.5 µM GAL-101.

**Figure 4:**
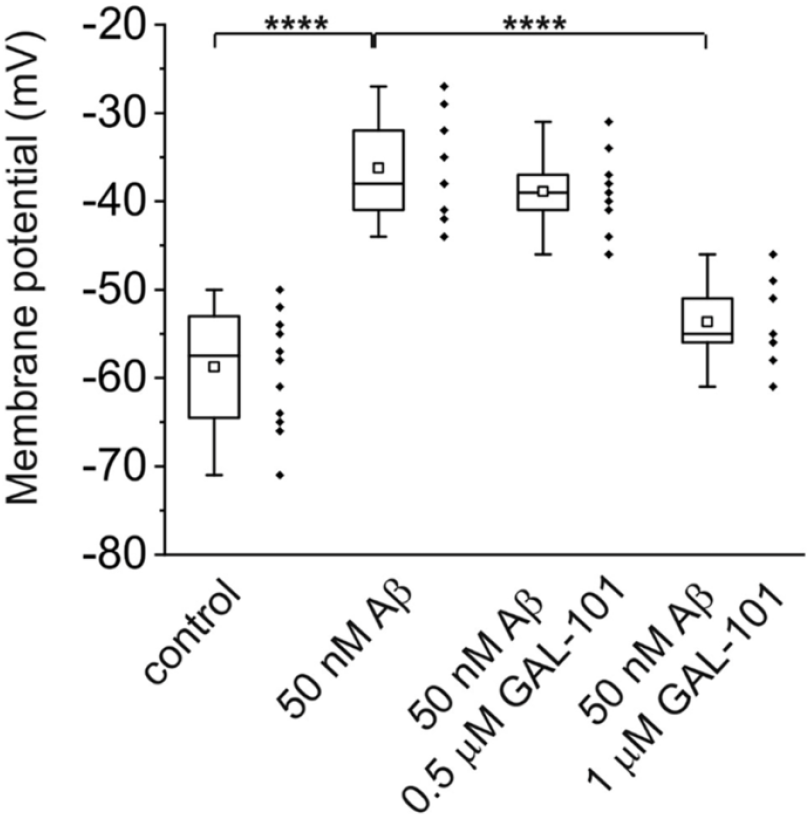
Chart plot of membrane potential measurements in control conditions (n=11), after addition of 50 nM Aβ (n = 8), and in the simultaneous presence of 50 nM Aβ and 0.5 μM (n = 9), or 1 μM GAL-101 (n = 9;p**** = 0.0001).

Figure 5A (shows the effect of 50 nM Aβ_1-42_ on the RMP of six RGC cells. After a variable time (1-15 minutes), the cells showed a membrane depolarization of several millivolts, reaching ∼ -20 mV. In figure 5B, six RGC cells were perfused with 50 nM Aβ_1-42_ in addition to 1 µM GAL-101. Even then, the RMP depolarized several minutes after the application of Aβ_1-42_ and GAL-101, reaching, however, less depolarized values (∼ -50 mV) than those recorded in the absence of 1 µM GAL-101. The time between Aβ_1-42_ perfusion and the onset of RMP depolarization was independent from the presence of GAL-101.

**Figure 5:**
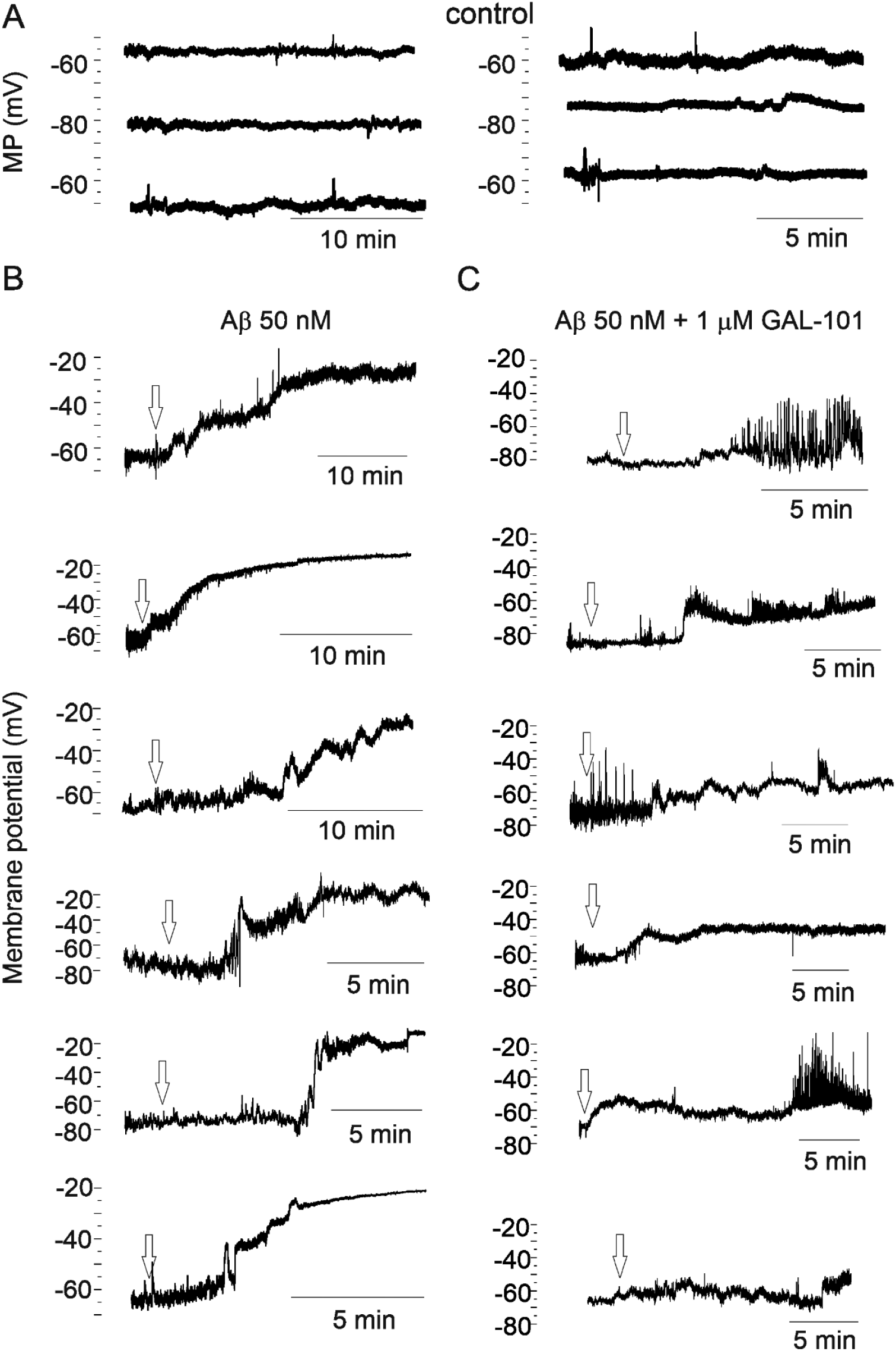
GAL-101 reduces Aβ_1-42_ oligomer-induced depolarization. Time-course current-clamp recordings of the individual RGCs’ resting membrane potential treated under control conditions (n = 6) (A), Aβ_1-42_ alone (n = 6) (B) and 50 nM Aβ_1-42_ together with 1 pM GAL-101 (n = 6) (C).

Figure 6 shows the RMP values under control conditions and after the application of Aβ_1-42_ alone or Aβ_1-42_ pre-incubated with 1 µM GAL-101. GAL-101 prevented major depolarization of RGCs.

**Figure 6:**
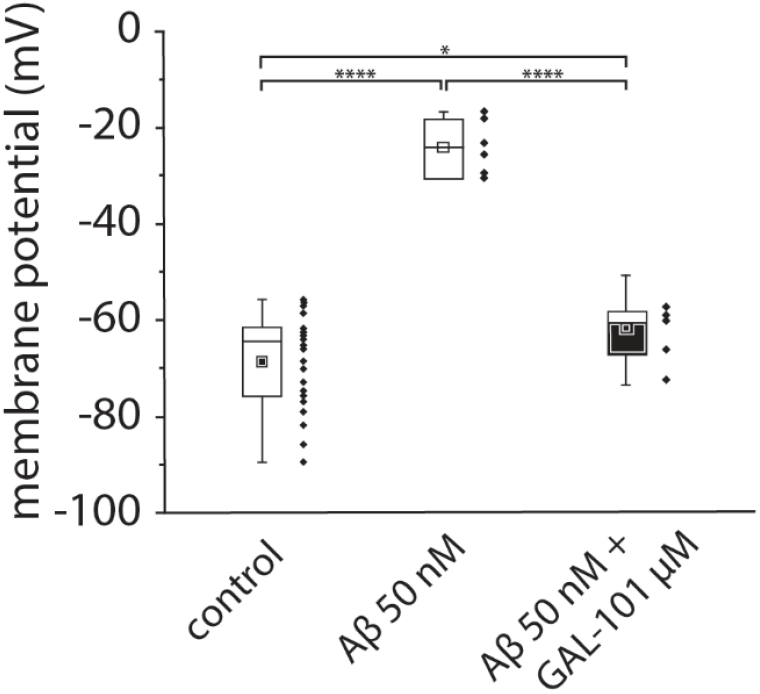
GAL-101 prevents depolarization of RGCs after 20 minutes. The chart plot shows the comparison between membrane potential values at the end of 20 minutes of membrane potential recording of single RGC cells in control conditions (n = 23), exposed to 50 nM Aβ (n = 6,p = 0.001) and 50 nMAβ+1 μM GAL-101 (n = 6, p = 0.001).

Since 50 nM Aβ_1-42_ significantly depolarizes the resting potential of RGCs, higher concentrations of Aβ_1-42_ could cause an inhibition of neuronal functions such as the initiation/conduction of action potentials and thus synaptic transmission. By preventing membrane depolarization, GAL-101 is expected to be able to counteract these negative effects.

### 3.3. GAL-101 effect on non-excitable cells

Long-term chronic depolarization is harmful for any kind of cell because most of the membrane transport systems are driven by the membrane potential. To test the ability of GAL-101 to interfere with Aβ_1-42_ effect in non-excitable cells, we performed electrophysiological recordings on RPE cells. Figure 7A shows the RMP values recorded from 29 RPE cells in current-clamp mode. Average RPE resting potential was -40.2 ± 2.4 mV. The same experiments were repeated, perfusing the cells with the extracellular solution containing the vehicle 0.1% DMSO. No differences were detected compared with control (data not shown). The difference between RGCs and RPE cells was the Aβ_1-42_ concentration required to induce a significant depolarization. The dose response/curve was obtained by incubating RPE cells at different Aβ_1-42_ concentrations and measuring the resting potential of the cells after 15 minutes (Figure 7B). From the best fit of the experimental points, it was possible to calculate an EC50 of Aβ_1-42_ for RPE cells (790 ± 80 nM) that was almost 18-fold less potent compared to the EC50 for RGCs (44.1 ± 6.3 nM). As with the RGCs, we used a concentration of Aβ_1-42_ able to depolarize the RMP without killing the cells. Figure 7C shows an example of the effect caused by 1 μM Aβ_1-42_ in a time course experiment in which the RMP was monitored for several minutes before and after the addition of Aβ_1-42_ (arrow). The effect of 1 μM Aβ_1-42_ under both control conditions and Aβ_1-42_ previously incubated with GAL-101 at two different concentrations is shown in Figure 8. GAL-101 at 10 times higher concentration than Aβ_1-42_ was not able to counteract the effect caused by 1 μM Aβ_1-42_ (Figure 8B; n = 7). However, GAL-101 at 20 times higher concentration than Aβ_1-42_ was able to inhibit the RMP depolarization seen in Figure 8A (Figure 8C; n = 11). As for RGCs, GAL-101 was able to significantly reduce the depolarization of the RPE cells but at a concentration 20-fold higher compared to Aβ_1-42_.

**Figure 7:**
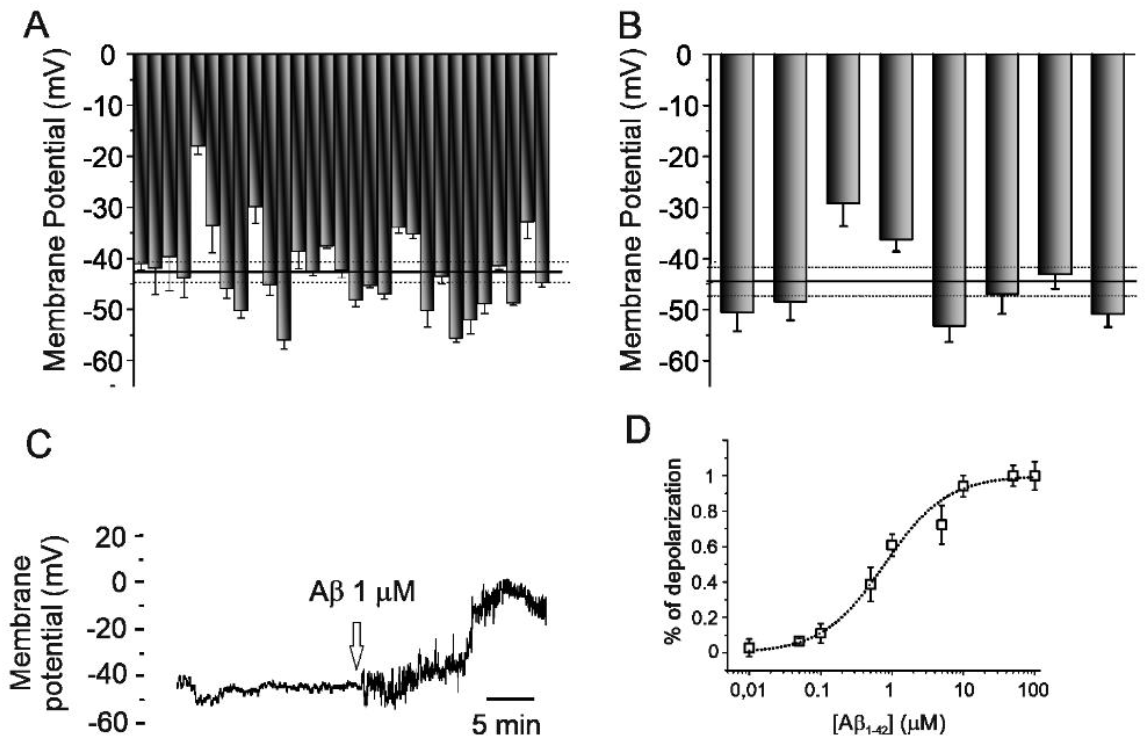
Aβ induces depolarization of retinal pigment epithelium (RPE) cells. A: Distribution of resting membrane values in RPE cells (n = 30). B: Distribution of resting membrane values in RPE cells perfused with the vehicle 0.1 % DMSO (n = 8). C: Time course experiment monitoring single RPE cell resting membrane potential during exposition to 1 μM Aβ (n = 5). D: Dose/response curve of RPE cells challenged with different Aβ concentrations EC50 (790 ÷ 80 nM, n = 8).

**Figure 8:**
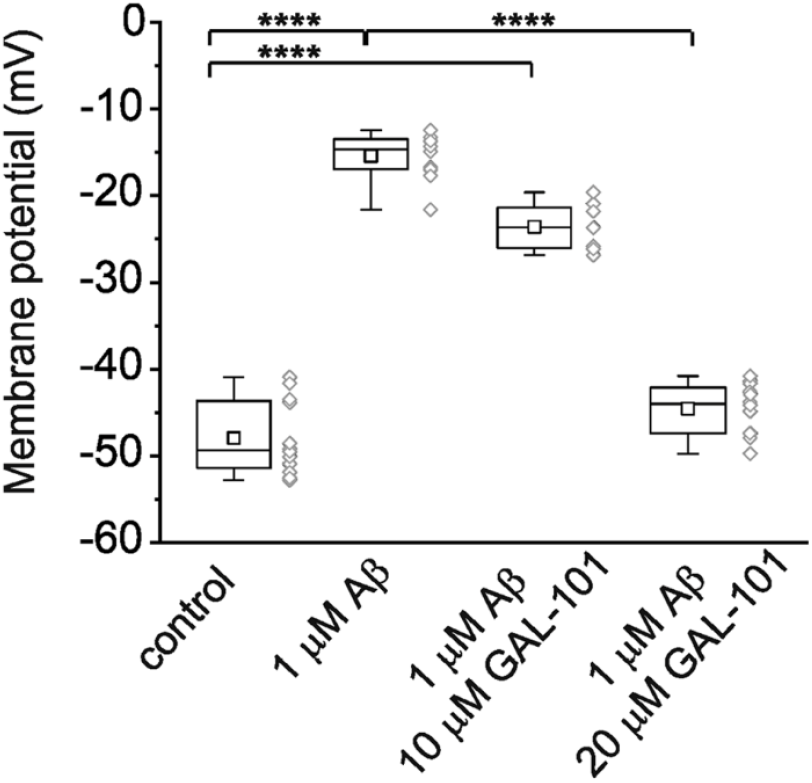
RPE depolarization induced by Aβ decrease in the presence of GAL-101. A: RPE resting membrane potential measurements under controlled conditions and in the presence of 1 μMAβ (−46.44 ±7.86 mV and -15.66 ±4.9 mV, respectively). B and C: Effect of GAL-101 at 10 μM (control vs. treated: -45.45 ± 5.43 mV and -23.6 ± 4.8 mV) or 20 pM (control vs. treated: -44.94 ± 5.69 mV and -41.56 ± 5.2 mV) respectively on RPE single cell depolarization induced by 1 μM Aβ. A andB:n=10;p**** = 0.0001.C:n= 10.

### 3.4. GAL-101 and amyloid ionic conductance in cell and artificial membranes

The ability of Aβ_1-42_ to operate as a transmembrane ionic pathway was tested in HEK cells under voltage-clamp, whole-cell configuration. The results presented in Figure 9, showed that 50 nM Aβ_1-42_ at 0 incubation time demonstrated the same conductance recorded once Aβ_1-42_ was incubated for 240 minutes (Figure 9, first and last open chart box from the left).

**Figure 9:**
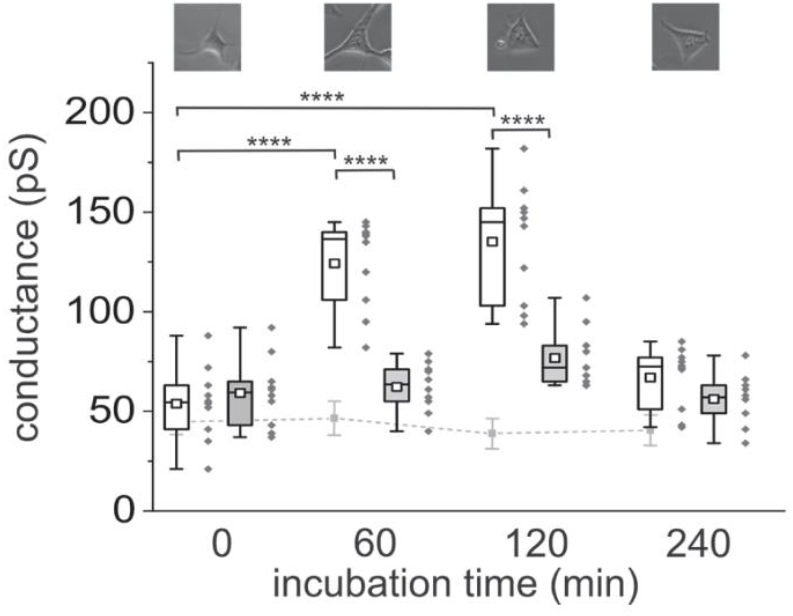
Membrane conductance is modified by Aβ_1-42_ Chart plot of whole cell HEK cell membrane conductance in the presence of 1 μM Aβ (white boxes) and with the addition of 20 μM GAL-101 (grey boxes). The gray line represents the experiments (n = 5at each time point) in the control condition (average and SE). 0 min: n= 10; 60 min: n= 10; 120 min: n =10; 240 min: n= 10.

However, Aβ_1-42_ maintained at 37 °C between 60 and 120 minutes and then added to the external solution caused a robust increase of HEK cells membrane conductance (Figure 9, second and third open chart boxes from the left). The same experiments were repeated with the Aβ_1-42_ previously incubated at 37 °C in the presence of 20 µM GAL-101, in accordance with the experiment presented in Figure 8. The final GAL-101 concentration in contact with the cells was then 200 nM in addition to 50 nM Aβ_1-42_. The membrane conductance measured under these conditions is plotted in Figure 9 at each incubation time as a grey filled chart box. In this case, the membrane conductance result was not significantly different under various conditions. Aβ_1-42_ preparation in the presence or in the absence of GAL-101 was also used to challenge an artificial bilayer. Tip-dip experiments resume, at single channel level, the results obtained in HEK cells. Figure 10 depicts current recordings after delivering Aβ_1-42_ after 120 and 240 minutes of incubation at 37 °C in the trans solution. The membrane conductance formed by the Aβ_1-42_ at different aggregation times show a marked difference in the current level between two- and four-hours incubation periods.

**Figure 10:**
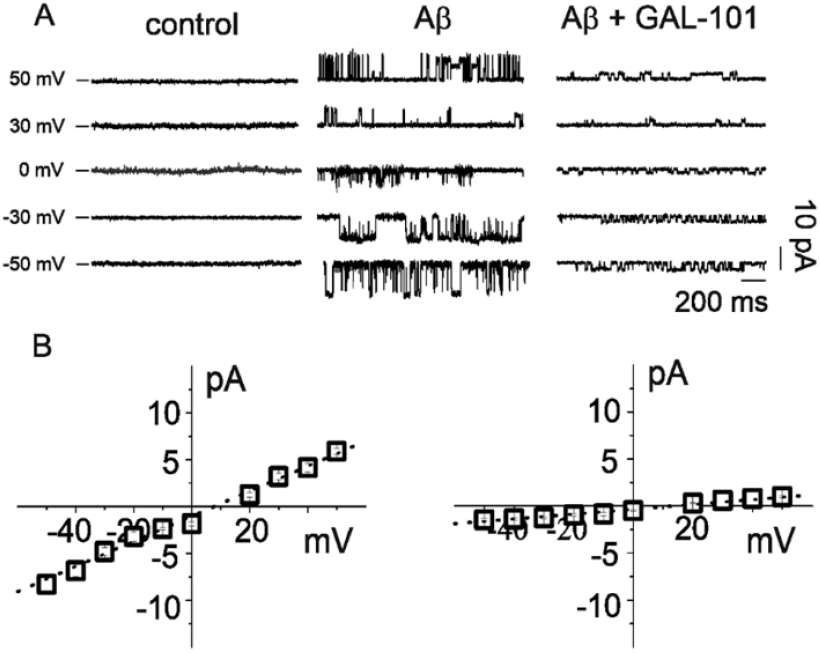
Tip-dip current recordings of Aβ_1-42_-induced membrane conductance in artificial lipid bilayer. A: Single channel recording at different membrane potentials under control conditions (left), in presence of 1 μM Aβ_1-42_ (incubated at37°C) in the absence (middle) and in the presence (right) of 20 μM GAL-101. B: Relative single channel current/voltage relationships. Control: n = 5; Aβ_1-42_ : n = 5; A _1-42_ + GAL-101: n = 5.

## 4. Discussion

Amyloid peptide aggregation and accumulation in the central nervous system (CNS) is a hallmark of several neurodegenerative diseases. In particular, Aβ_1-42_ concentration in the CNS fluid is one of the tools used to identify patients with AD, and it is combined with other biomarkers such as phosphorylated tau and advanced imaging techniques to get a more accurate AD diagnosis (40,41). Several reports have shown that Aβ_1-42_ deposits are also present in different retinal cell layers and in the optic nerve (15,16). Both the presence of the peptide and several degenerative processes in the eye have been proposed as biomarkers to identify neurodegenerative pathologies (42). For example, it is not unusual that AD patients also suffer from degeneration of the retina (26,43). Glaucoma and AMD are the most recognized causes of irreversible vision loss worldwide. Indeed, both these pathological conditions show Aβ_1-42_ deposits in the retina (44,45).

Although the molecular mechanism by which Aβ_1-42_ exerts its cytotoxicity has still not been completely elucidated, it is widely accepted that the toxic form of the amyloid peptide is represented by the soluble oligomeric aggregates (46–49). The neurodegenerative action of Aβ_1-42_ oligomers is believed to be exerted both at the membrane level (50,51) as well as inside the cell after cytoplasmic inclusion of Aβ_1-42_ agglomerates (52–54).

Several membrane receptors have been identified to react to Aβ_1-42_, either by increasing their activity or inhibiting their functions (55–60). It is still controversial if Aβ_1-42_ can directly cause membrane instability and/or have the ability to form ion-conducting channels in vivo. Certainly this is true for artificial lipid bilayers where Aβ_1-42_ is responsible for generating conductance directly proportional to the Aβ_1-42_ aggregation state (35).

Despite not evaluating the exact mechanism of action of Aβ_1-42_, in these experiments, we demonstrated that Aβ_1-42_ oligomer-enriched solution at 50 nM concentration was able to cause RMP depolarization of isolated mouse RGCs. According to the cell properties, opening of non-selective ionic conductance should move the cell resting potential to more depolarized values. This is exactly what happened in our experiments, suggesting the formation of ionic conductance more than overt, acute membrane damage. Furthermore, the kinetics of the change in the membrane potential was a relatively slow process, suggesting a step-by-step addition of small ionic pathways, more than the rapid acute response of e.g., ionic membrane receptors. However, we cannot exclude the partial or total contribution of a slowly activating receptor such as a G protein-coupled receptors (GPCR) in causing membrane depolarization.

In light of these results, a strategy to reduce RGC depolarization could be to block Aβ_1-42_ aggregation to prevent the formation of the toxic Aβ_1-42_ oligomers. For GAL-101, it was demonstrated that this compound prevents and even reverses the formation of Aβ_1-42_ oligomers (61). The results on RGCs demonstrated that GAL-101 is able to prevent amyloid-induced GC depolarization, possibly acting on amyloid aggregation formation. Since the effect of Aβ_1-42_ was the depolarization of membrane potential, the final outcome would be the inhibition of neuronal functions such as the generation and conduction of action potentials and synaptic transmission. The antiaggregant action of GAL-101 fully prevents the depolarization of the RMP, thus preserving the neuronal activity of RGCs.

Aβ_1-42_-induced membrane potential depolarization was also effective on non-excitable cells as shown by RPE recordings. However, the concentration of Aβ_1-42_able to cause a change in the RMP was 20-fold higher for RPE cells compared to RGCs. A 20-fold difference in the EC50 of Aβ_1-42_ as determined in RPE cells and in RGCs indicates that the mechanism of toxic action is different in these two cell types: the first has epithelial origin and the second has neuronal origin. The toxicity to the neuronal RGCs might be caused by an interaction of Aβ_1-42_ with one of the described binding sites located on neuronal receptors, e.g., glutamatergic, GABAergic, or nicotinic receptors (49), while the toxicity to the epithelial RPE cells might be caused by a lower affinity interaction with the membrane. The toxic effect of Aβ_1-42_ on the neuronal RGCs was very strong, with an EC50 of 44 nM calculated from the dose/response curve (Figure 3C). RPE cells required a much higher Aβ_1-42_ concentration able to induce a significant depolarization (Figure 6C). It is possible that the lipid membrane composition is different between these two cell types, making the insertion of Aβ_1-42_ aggregates into the cells’ plasma membrane more difficult. A second possibility is that RGCs express a higher number of membrane receptors than RPE cells able to react with Aβ_1-42_, thus contributing to the cell depolarization. Both hypotheses are supported by the similar action of GAL-101 on the two cell types. Our experiments show that this anti-aggregation compound must be 20 times more concentrated than Aβ_1-42_ to exert its activity.

In conclusion, we have shown that Aβ_1-42_ oligomers are able to cause membrane potential depolarization both in neuronal and epithelial cells from retinal tissue. We suggest that the mechanism of action is the insertion of oligomeric Aβ_1-42_ in the cell membrane and the opening of non-selective ionic permeability pathways. In addition to this non-specific effect, it is possible that RGCs expressing membrane receptor are able to react with Aβ_1-42_, contributing to membrane depolarization. Evidence that the RMP in the presence of Aβ_1-42_ does not achieve 0 mV strongly suggests that the change in the membrane permeability is not due to membrane disruption. With membrane breaks, in a very short time, the cytoplasm would wash out by the external solution. On the contrary, the activation of membrane receptors or, more likely, the formation of non-selective pores, for the physiological characteristics of the cell, stabilizes the membrane resting potential to -10/-15 mV. Pre-incubation of Aβ_1-42_ during the aggregation process with GAL-101, at the concentration tested in our experiments, inhibits amyloid peptide aggregation. This should prevent strong changes in resting membrane potential and thus would prevent impairment of cell function.

A possible future therapeutic option could be the chronic delivery of GAL-101. After treatment, Aβ_1-42_ aggregation process should slow down and, consequently, this would eventually slow down membrane depolarization during various retinal pathological stages. A potential treatment approach for glaucoma and dry AMD, which are caused by the neurodegeneration of RGCs and RPE cells, could be based on the elimination of Aβ_1-42_ oligomers from the retina that have detrimental effects on the RMP of these cells, leading to long-term toxic effects.

A potential drug candidate for this approach could be the Aβ_1-42_ aggregation modulator GAL-101, which has already shown beneficial effects in an animal glaucoma model (62). The prevention of Aβ_1-42_-induced depolarization in both RGCs and RPE cells, together with the known effects of GAL-101 on Aβ-suppressed synaptic plasticity (61,63), qualifies GAL-101 as a new candidate for further clinical investigation in Aβ-associated retinopathies.

## 6. Author Contributions

Individual contributions. “Conceptualization, C. P. and M.M.; methodology, E.P., S.G and S.S.; validation, D.B., C.P. and M. M.; formal analysis, E.P, S.G., C.A.M; investigation, C.P. and M.M.; resources, M.M.; data curation, S.S and D.B.; writing—original draft preparation, C.P., E.P., S.G., S.S., and D.B.; writing—review and editing, C.A.M., H.R., M.M.. All authors have read and agreed to the published version of the manuscript.”

## 8. Acknowledgments

This manuscript is dedicated to the late Christopher G. Parsons, an exceptional pharmacologist who made a significant contribution to this scientific work with his passion for the fascinating mechanism of action of GAL-101. He was our friend and colleague, while always reminding us that the science behind GAL-101 is extraordinary and well worth the extra effort to advance it.

